# HPDL is critical in human cortical development via regulation of mitochondrial functional properties

**DOI:** 10.1101/2025.04.28.651017

**Authors:** Matteo Baggiani, Maria Andrea Desbats, Valentina Naef, Michela Giacich, Daniele Galatolo, Serena Mero, Sara Zampieri, Valentina Cappello, Agata Valentino, Leonardo Salviati, Filippo Maria Santorelli, Devid Damiani

## Abstract

Human brain development is highly regulated by several spatiotemporal processes, which disruption can result in severe neurological disorders. Emerging evidence highlights the pivotal role of mitochondrial function as one of these fundamental pathways involved in neurodevelopment. Our study investigates the role of 4-hydroxyphenylpyruvate dioxygenase-like (HPDL) protein in cortical neurogenesis and mitochondrial activity, since mutations in the *HPDL* gene are associated with SPG83, a childhood-onset form of hereditary spastic paraplegia characterized by corticospinal tract degeneration and cortical abnormalities. Starting from mutant neuroblastoma cells, we demonstrated that HPDL is essential to mitochondrial respiratory chain supercomplex assembly and cellular redox balance. Moreover, transcriptomic analyses revealed dysregulated pathways related to neurogenesis, implicating HPDL role in early cortical development.

To further elucidate the role of HPDL, we generated cortical neurons and organoids from SPG83 patient-derived induced pluripotent stem cells. Mutant cells exhibited premature neurogenesis at early differentiation stages, likely leading to depletion of cortical progenitors, as evidenced by decreased proliferation, slight increase of apoptosis, and unbalanced cortical type composition at later stages. Furthermore, cortical organoids derived from SPG83 patients showed impaired growth, reminding microcephaly observed in severe cases. In addition, mitochondrial morpho-functional characterization in mutant neurons confirmed disruption of OxPhos chain functionality and increased ROS generation rate. Treatment of cortical cells with two antioxidant compounds, could partially revert premature neurogenesis.

In conclusion, our findings reveal a critical role for HPDL in coordinating cortical progenitor proliferation, neurogenesis, and mitochondrial function. These insights shed light on a mechanistical understanding of SPG83 pathology and underscore the therapeutic potential of targeting oxidative stress in this and related neurological disorders.

## Introduction

Hereditary Spastic Paraplegias (HSPs) cover a family of heterogeneous rare genetic diseases of corticospinal tract structure and function and presenting as pure or complex forms where paraparesis is accompanied by additional neurological manifestations, such as peripheral neuropathy, second motor neuron involvement, and cerebellar ataxia.^1,2^ HSPs have a complex genetic etiology with approximately 90 different genes identified to date.^3,4^ Mutations in the *HPDL* gene, encoding the protein 4-<u>H</u>ydroxy<u>P</u>henylpyruvate <u>D</u>ioxygenase-<u>L</u>ike, have been associated with SPG83 (<u>SP</u>astic paraple<u>G</u>ia 83), documented in over 90 individuals from approximately 50 families worldwide.^5^ The spectrum of symptoms ranges from neurodevelopmental disorders, including cerebral palsy, epilepsy, and other features resembling Leigh syndrome, white matter abnormalities, and mild adolescent-onset forms of progressive spasticity in the lower limbs.^6–12^ *HPDL* is the only known mammalian paralogue of the *HPD* gene, which is involved in tyrosine metabolism, though its function appears to be different. HPDL is characterized by two vicinal oxygen chelate domains and three iron-binding sites, mitochondrial localization thanks to its N-terminus mito-targeting sequence, and high expression in the brain. Despite the rapid identification of individuals carrying biallelic *HPDL* mutations, the specific biological function of HPDL remains largely unknown. In *vivo* studies exploring HPDL-related conditions have faced several limitations. In particular, *Hpdl*^-/-^ mice exhibited early lethality and reduced brain sizes with cortical apoptosis and epilepsy,^7^ reminiscent of the features observed in a subset of SPG83 children. Similarly, zebrafish *hpdl* morphants showed motor response impairments.^11^ Unfortunately, both the transient nature of the fish model and the perinatal lethality in mice prevent longitudinal work. Moreover, studies in non-neural cell lines did not reveal significant mitochondrial dysfunction, whereas neural cell lines and brain tissues exhibited impaired cellular respiration.^7^ Recent investigations based on oxygen-based metabolome suggested that HPDL could convert the tyrosine catabolite 4-hydroxyphenylpyruvate to 4-hydroxymandelate, being involved in an alternative pathway for the biosynthesis of coenzyme Q_10_ (CoQ_10_).^13^ This role could represent a key step, considering that CoQ_10_ biosynthesis deficiencies are associated with a growing rate of neurological conditions,^14^ and ubiquinone supplementation in a group of SPG83 children holds promise in improving in part their clinical manifestations (Santorelli FM, unpublished observation).

The core hypothesis of this work is the crucial role that HPDL could play in neural and brain development. We conducted *in vitro* experiments using HPDL knockout (KO) in neuroblastoma cell lines (SH-SY5Y) as well as bi- and tridimensional cortical tissues differentiated from SPG83 patient-derived induced pluripotent stem cells (iPSCs).^15^ We have recently generated and validated for cortical specification and neuronal maturation.^16^ In this work we finely characterized the effect of *HPDL* variants during early cortical differentiation (about one month) and reported the critical role of HPDL for regulating the balance between proliferation and neurogenesis of cortical progenitors. Additionally, we observed that HPDL KO SH-SY5Y and HPDL mutant cortical cultures exhibited mitochondrial dysfunction, increase of reactive oxygen species (ROS), and apoptosis. We also showed that treatment with antioxidants such as MitoTEMPO can partially rescue anticipation of neurogenesis, posing the basis for future pharmacological approaches in clinical practice.

## Materials and Methods

### Cell Reprogramming

Human dermal fibroblasts (HDFs), obtained from patient skin tissue via punch biopsies, were amplified in HDF medium and reprogrammed following a modified form of an established protocol,^17^ as previously described.^18^ Briefly, dissociated HDF cell suspension were transfected via Amaxa Nucleofector 2b (Lonza) with a mix of episomes for integration-free expression of human “Yamanaka’s factors” (L-MYC, LIN28, OCT3/4, SOX2, and KLF4). Transfected HDFs (tHDFs) were cultured on Geltrex-coated dishes (A1413301, Thermo Fisher Scientific) with E7 medium, 0.5 mM sodium butyrate (B5887, Sigma), 1 μM hydrocortisone (H0888, Sigma), checking positive expression of mCherry after 2 days.^19^ From about 14 days onwards, iPSC “islands” appeared, and medium was switched to StemFlex medium (A3349401, Thermo Fisher Scientific). iPSC clones with uniform flat and round shaped morphology were picked in sterile conditions and propagated as single cell lines. After seven passages, all iPSC clones were characterized as previously described.^16,18^

### Cell culture

Neuroblastoma SH-SY5Y cells, iPSCs, neural progenitors, and neurons were cultured in standard conditions at 37 °C and 5% CO_2_. SH-SY5Y were maintained in DMEM High Glucose (ECM0728L, Euroclone) containing 15% FBS (SIAL-FBS-SA; Sial), 1% MEM-NEAA (ECB3054D, Euroclone) and 1% Pen/Strep (SIAL-PEN/STREP, Sial). To prompt metabolic switching from glycolysis to respiration without affecting cell viability SH-SY5Y cells were grown for 48 h in DMEM/F12 no glucose (PM150322, Elabscience), supplemented with 2 mM D-Glucose (16325, Riedel-de Haen), 15% FBS, 2mM L-glutamine (25030081, Thermo Fisher Scientific) and 100 µg/ml Pen/Strep (OxPhos medium).

Human iPSCs were cultured on Geltrex coated 6-well plates in StemFlex medium, refreshing medium every other day. Cells were passed every 5-7 days with ReLeSR™ (100-0484, Stem Cell Technologies), following manufacturer instructions. Cells have been differentiated following the dual-SMAD inhibition protocol.^20,21^ Briefly, pluripotent cells were harvested at high confluence on Geltrex and cultured for 12 days in neural induction medium containing 100 nM LDN193189, 10 µM SB431542, and 2 µM XAV939 (72147, 100-1051, 72674 respectively, Stem Cell Technologies), splitted with Accutase (A1110501, Thermo Fisher Scientific) and replated on poly-D-lysine/laminin coated coverslips (A3890401, Thermo Fisher Scientific and L2020, Sigma, respectively). From day 16 onward, medium was switched to terminal differentiation medium containing 30 ng/ml BDNF (AF-450-02, PeproTech) to improve neuronal differentiation and maturation. Differentiated cells were harvested at day *in vitro* (DIV) 16 and DIV30 and processed for immunostaining (see below) or RNA extraction via RNeasy Plus kit (74134, Qiagen).

Concerning the treatment with antioxidants, 1 mM glutathione monoethyl ester (GSH-MEE; 353905, Merck) or 1 µM MitoTEMPO (SML0737, Merck) were added on neural progenitors at DIV12 and DIV14.^22–24^ The cells were fixed at DIV16 after a 4-days treatment.

### Organoid culture

Cortical organoids were generated with a slight modification of the classical SFEBq-based procedure.^25^ Briefly, 9.000 iPSCs for well were plated in 96-well plates (83.3925.400, Sarstedt) in StemFlex medium with Y-27632 10 µM (ab120129, Abcam). The day after, we performed one wash with PBS and added organoid medium I, containing Glasgow’s MEM (11710035, Thermo Fisher Scientific), 1% Pen/Strep, 20% KnockOut Serum Replacement (10828010, Thermo Fisher Scientific), 1% MEM-NEAA, 1mM Pyruvate (S8636, Sigma), 0.1 mM 2-mercaptoethanol (21985, Gibco), for 18 days, changing the medium twice a week. Cell aggregates were then transferred on ultra-low-adherent 24-well plates (662102, Greiner) in organoid medium II, containing DMEM/F12 – Glutamax (10565018, Thermo Fisher Scientific), 1% N2 (17502001, Thermo Fisher Scientific), 1% Chemically Defined Lipid Concentrate (11905031, Thermo Fisher Scientific), 1% Pen/Strep, until 35^th^ day. Medium was changed once a week. For size quantification, organoids were imaged with Zeiss Axiovert 25 at DIV5 and Leica M205 FA at DIV10 and 35.

### Cytochemical staining for cytochrome-c oxidase and succinate dehydrogenase activity

For cytochemical staining for the enzymatic activities of cytochrome-c oxidase (COX) and succinate dehydrogenase (SDH), neurons were processed using standard protocols.^26^ Briefly, cells were incubated for one hour at RT with a solution containing phosphate buffer (0.1 M, pH 7.6), 75 mg/ml sucrose (S7903, Merck), 1 mg/ml cytochrome-c (C7752, Merck), 2 μg/ml catalase (C40, Sigma), and 0.5 mg/ml DAB (D5637, Merck). For SDH staining, cells were incubated instead for 1 hour at 37 °C in a phosphate buffer medium (0.2 M, pH 7.6) containing 27 mg/ml sodium succinate (S2378, Merck) and 1 mg/ml Nitro Blue Tetrazolium (N6876, Merck). After incubation, cells were rinsed twice with phosphate buffer, fixed 15 minutes with 4% paraformaldehyde in PBS, washed twice with PBS and then imaged with Zeiss Axiovert 25 microscope.

### DCFDA staining

DCFDA (C6827, Invitrogen) assays were performed on neural progenitors or neurons following a single protocol. After removing the medium, the cells were washed once with H-HBS solution (HBSS (14175095, Thermo Fisher Scientific) containing HEPES (EMR152100, Euroclone) 1 mM pH 7.4). DCFDA 1 µM and Hoechst 33342 5 μg/ml were diluted in H-HBS solution and added on cells for 40 min at 37°C in darkness. Finally, one wash in H-HBS solution was performed and the cells were acquired with Zeiss LSM 900 confocal microscope.

### Fluorescence quantification

Fiji software was used to analyze the mean fluorescence of TMRM, DCFDA, Nestin or TUBB3. In particular, images were thresholded keeping the values constant inside every differentiation and used to select ROIs. Then, the mean fluorescence of selected marker was measured inside the ROI. Every measure was normalized on CTRL line of every differentiation.

### Assays of Respiratory Chain Enzyme Activities

Prior to the measurements, cells were cultured for 48 h in OxPhos medium. Activities of mitochondrial respiratory chain complexes I, II, III, IV, and II+III were measured as previously described^27^ and normalized to protein and citrate synthase (CS) activity using the Varian Cary 100Bio UV-Visible Spectrophotometer.

### Statistical analysis

All data in the manuscript represent three or more independent experiments. We performed the statistical analysis using GraphPad Prism 9.0.0 software. All data were analyzed using either parametric or non-parametric methods, based on the normal distribution analysis. The significance among groups with 2 independent variables was determined using two-way ANOVA test, among 3 or more groups with 1 independent variable was measured with one-way ANOVA or Kruskal-Wallis tests, and between 2 groups with 1 independent variable was calculated with two-tailed unpaired Student’s t-test. Statistical significance is reported as: * *P*-value ≤ 0.05, ** *P*-value ≤ 0.01, *** *P*- value ≤ 0.001, or **** *P*-value ≤ 0.0001.

### Ethical Statements

The study was conducted in accordance with the 333 Declaration of Helsinki and approved by the Ethics Committee of Tuscany Region CEPR (protocol 334 code 102/2020, date of approval 05/05/2020). Written informed consent was obtained from all subjects involved in the study.

### Data availability

Data will be publicly available upon request.

## Results

### Neural derived HPDL KO cell lines display mitochondrial functional impairment

To delve into the molecular function of HPDL in SPG83 etiopathogenesis, we generated a HPDL KO SH-SY5Y cell line (c.318delC; p.Val107PhefsTer24) via CRISPR/Cas9 gene targeting (**Fig. 1A**). The mutation obtained resulted in absence of full length HPDL protein as shown by Western Blot (WB) analysis, whereas specific signals were present in cells transfected with empty vector (lacking any specific sgRNA; from now on, E.V. cells) or wild type (WT) SH-SY5Y cells (**Fig. 1B**). Given the known mitochondrial localization of HPDL, we assessed mitochondrial activity by analyzing the expression levels and assembly state of the respiratory chain by Blue Native-PAGE (BN-PAGE). Formation and/or stability of mitochondrial respiratory chain supercomplexes (RCSs) containing complexes I, III_2_, and IV, dimeric complex V, and individual complexes III, IV, and V, appeared reduced in KO cells (**Fig. 1C, Supplementary Fig. 1A, B**), with normal levels of complex II. The reduction in complexes I and IV seems to be partly explained by significant decreased levels of NDUFB8 and COXII (subunits of complexes I and IV, respectively) in KO compared to E.V. cells, while levels of complexes II (SDHA), III (UQCRC2), and V (ATP5A) subunits remained unchanged, excluding a general decrease of mitochondrial proteins expression in HPDL KO cells (**Fig. 1D**). Assessment of oxidative metabolism in KO cells showed a reduction in respiratory chain enzyme activities of complexes I, III, and IV (but not complexes II and II+III; **Fig. 1E**), whereas micro-oxygraphy showed impaired oxygen consumption with reduced basal and ATP-coupled respiration rates (**Fig. 1F**) and unaltered maximal and spare respiratory capacities in HPDL KO cells (**Fig. 1F, Supplementary Fig. 1C**). In spite of this, the levels of total CoQ_10_ increased in KO cells (**Supplementary Fig. 1D**), whereas copy number of mtDNA was similar in E.V. and HPDL KO cells (**Supplementary Fig. 1E**). In line with these results, production of ROS appeared increased in stressed KO cell line (**Fig. 1G**). Altogether, these results indicate that HPDL ablation impaired OxPhos metabolism.

**Figure 1.**
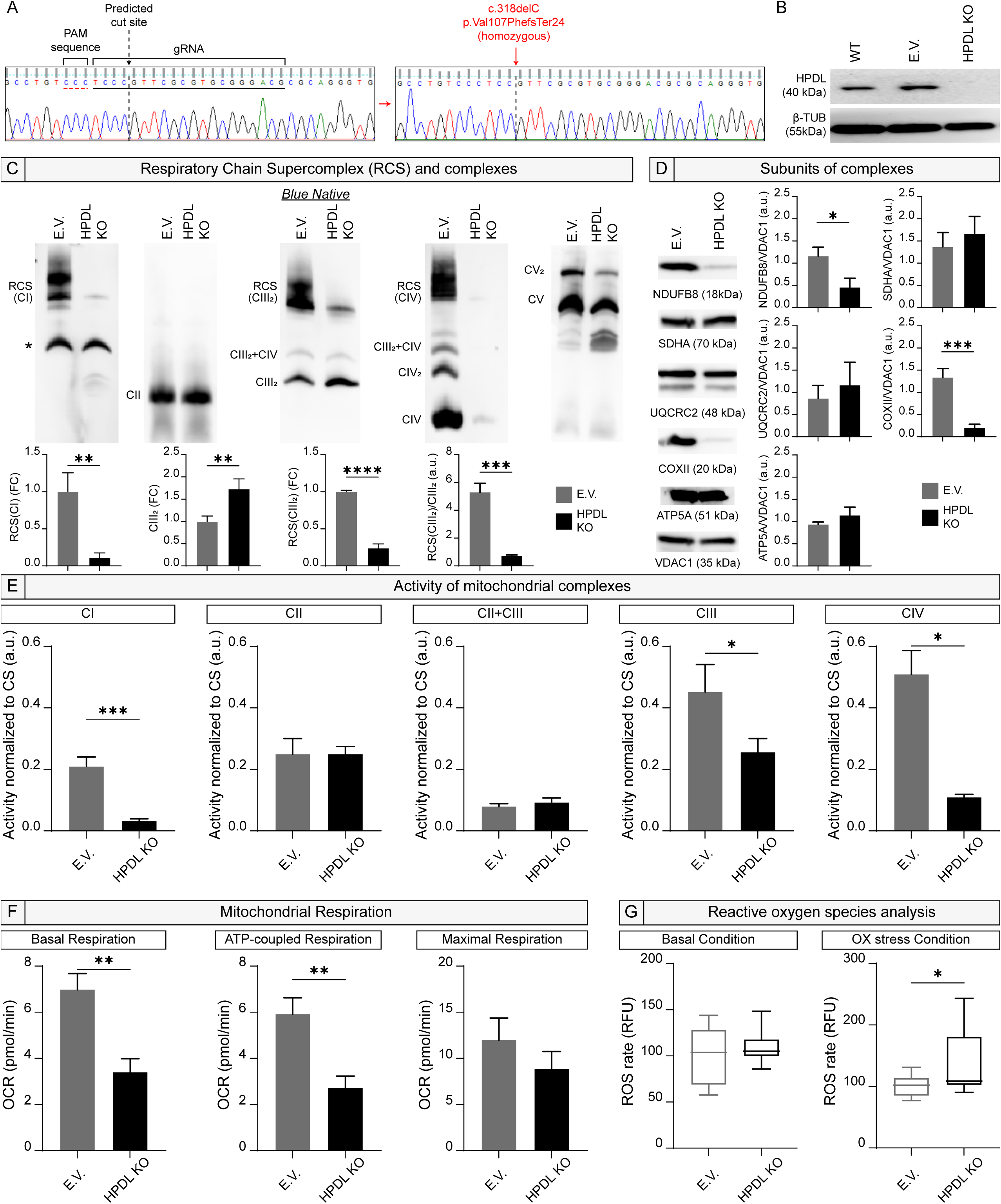
Generation of HPDL KO SH-SY5Y cell line and biochemical analysis. (**A**) Electropherogram of predicted cut site using CRISPR-CAS9 technique and electropherogram showing c.318delC mutation in HPDL sequence after cutting. (**B**) WB analysis shows the presence of full length HPDL protein in WT and E.V. SH-SY5Y cell lines and absence in HPDL KO SH- SY5Y cells. (**C**) BN-PAGE images exhibit the decrease in the levels of RCSs, individual complexes III_2_ (CIII_2_) and IV (CIV), and dimeric complex V, while no changes are found in complex II (CII). Bar plots indicate the significant decrease of RCS(CI), RCS(CIII_2_), and RCS(CIII_2_)/CIII_2_ (unpaired Student’s t-test, *n* = 3, *P* = 0.0043, unpaired Student’s t-test, *n* = 3, *p* < 0.0001, unpaired Student’s t-test with Welch’s correction, *n* = 3, *P* = 0.0061) and the proportional increase of CIII_2_ (unpaired Student’s t-test, *n* = 3, *P* = 0.0081) in HPDL KO SH-SY5Y cells compared to E.V. ones. All data in bar plots are represented as mean ± SD. (**D**) Gel images of loading control and corresponding bar plots show the significant reduction of NDUFB8 and COXII (*n* = 3, *P* = 0.0130 and 0.0008, respectively) and no changes in the levels of SDHA, UQCRC2, and ATP5A (*n* = 3, *P* = 0.3615, 0.4346, and 0.1346, respectively). All data in bar plots are normalized to the mitochondrial protein VDAC1 and represented as mean ± SD. All statistical analyses were conducted using unpaired Student’s t-test. (**E**) Bar plots show the decrease of activity of mitochondrial CI, CIII_2_ (unpaired Student’s t-test, *n* = 3, *P* = 0.0006 and 0.0274, respectively), and CIV (unpaired Student’s t-test with Welch’s correction, *n* = 3, *P* = 0.0115) in HPDL KO SH-SY5Y line compared to E.V. one, normalized to Citrate Synthase (CS). CII and CII+CIII_2_ mitochondrial activities do not exhibit any significant changes among lines (unpaired Student’s t-test, *n* = 3, *P* > 0.9999 and 0.2746, respectively). All data are represented as mean ± SD. (**F**) Oxygen Consumption rate (OCR) measured via Seahorse XF analysis demonstrates a decrease in Basal and ATP-coupled respiration in HPDL KO cells compared to E.V. ones (unpaired Student’s t-test, *n* = 3, *P* = 0.0025 and 0.0032, respectively), while no difference among them is found in Maximal Respiration (unpaired Student’s t-test, *n* = 3, *P* = 0.1508). All data in bar plots are represented as mean ± SD. (**G**) Plots show comparison of ROS species in basal and oxidative (OX) stress conditions, demonstrating an increase in OX stress conditions in HPDL KO compared to E.V. lines (unpaired Student’s t-test, *n* = 3, *p* = 0.3072 and unpaired Student’s t-test with Welch’s correction, *n* = 3, *P* = 0.0392, respectively). All data are represented as box and whisker plot, min-to-max.

To clarify how loss of function of HPDL impacts on gene expression, we performed bulky RNA-seq analysis in E.V. and mutant cells (**Supplementary Fig. 1F**). Results showed a set of 3052 differentially expressed genes (DEGs) of which 1438 were upregulated and 1614 downregulated (**Supplementary Fig. 1F, G**). Interestingly, profound effect was exerted on metabolism, with significant enrichment in genes related to oxidative metabolism such as response to oxygen levels, reactive oxygen species biosynthetic process, and NAD metabolic processes (**Supplementary Fig. 1G**). In addition, several genes involved in extracellular matrix organization appeared to be upregulated, whereas genes working in *Signal Release from Synapse*, *Neurotransmitter Secretion*, and function were mostly downregulated (**Supplementary Fig. 2A**). Gene Ontology (GO) analysis defined that most of DEGs belonged to categories related to brain development, structure, and function, such as *Synapse Organization and Assembly*, *Positive Regulation of Neuron Differentiation*, *Regulation of Membrane Potential*, *Axon and Neuron Projection Guidance*, *Neurotransmitter Transport,* and *Secretion* (**Supplementary Fig. 2B**).

### Anticipation of neurogenesis in SPG83 patient-derived iPSC lines

The results obtained from the neuroblastoma SH-SY5Y cell line prompted us to explore the role of HPDL in cellular models that more closely resemble the brain dysfunction occurring in SPG83. To this end, we compared immunocytochemical and transcriptomic profiles of cerebral cortical tissue differentiated from iPSC lines deriving from two healthy donors and four SPG83 patients carrying different mutations in the *HPDL* gene. We previously described generation of SPG83 iPSCs from Patient 1 (carrying the homozygous missense p.Ser49Arg variant), Patient 2 (harboring a missense p.Ile266Thr variant in compound heterozygosity with a frameshift p.Ala86ArgfsTer45), and Patient 3 (carrying a compound heterozygote p.Phe31Leu and p.Gly278Ser missense^16^). Here, we also used a fourth iPSC line derived from a 12-year-old girl (Patient 4) harboring the homozygous p.Gly50Asp HPDL variant, as well as the commonly used ACS-1019 iPSC line (ATCC), from now on Control-1 or CTRL-1), and a control line derived from a 37-year-old healthy female donor (from now on, Control-2 or CTRL-2). Fine characterization of these two iPSC lines is reported in **Supplementary** Fig. 3.

Control and HPDL mutant cells were differentiated in dorsal telencephalon following established protocols,^20,21^ known to produce cortical neural progenitors after 16 days of differentiation (DIV16). Corroborating our earlier work, we observed that HPDL mutant cells at DIV16 were correctly differentiated in anterior neural tissue, as the totality of neural cells displayed homogeneous positivity for the telencephalic marker FOXG1 (**Fig. 2A**).

**Figure 2.**
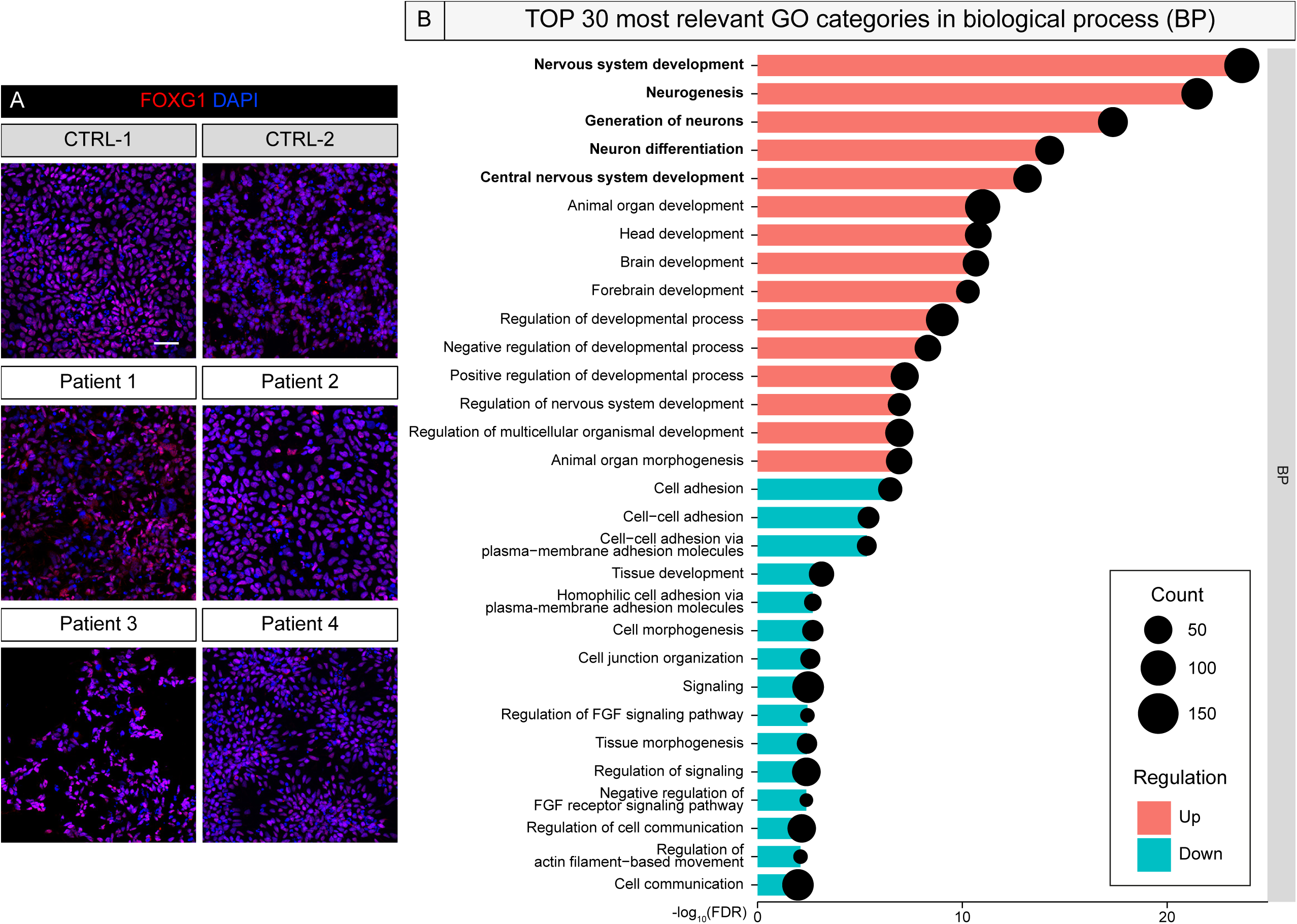
HPDL protein in FOXG1 positive cortical neural progenitors and Gene Ontology analysis. (**A**) Representative confocal images of telencephalic marker FOXG1 in CTRL and HPDL neural progenitor cells (DIV16). Scale bars are 20 μm. (**B**) GO analysis of DEGs in HPDL vs Control neural progenitor cells (DIV16) showing the TOP 30 categories in biological processes (BP), sorted for False Discovery Rate (FDR), and the number of gene counts. All nuclei are stained with DAPI.

Then, we compared control vs HPDL mutant progenitors at the same DIV16 time point by RNA-Seq. GO analysis performed on 785 DEGs (492 up- and 293 downregulated) implied deregulation of fundamental pathways involved in neurodevelopment, with genes clustering in categories such as *Forebrain Development*, *Neuron Differentiation*, and *Cell-Cell Signaling*, as already observed in HPDL KO SH-SY5Y cells. Notably, GO categories related to regulation of neurogenesis were the most significant in the analysis (**Fig. 2B**) with evident upregulation of neurogenesis-related genes such as *NEUROG2*, *NEUROG3*, and *BCL6* (**Fig. 3A, Supplementary Fig. 4A**). Genes related to cortical development (like *OTX1*, *FOXP2*, and *LMX1A/B*) and WNT pathway (such as *WNT3A*, *WNT7A*, *WNT11*, *LEF1*, *FZD7*, *AXIN2*), known to increase expression and operate fundamental regulation in the process of cortical neurogenesis,^28–30^ also appeared to be upregulated, while genes related to FGF pathway, more linked to proliferative state,^31^ seemed to be mostly downregulated (**Fig. 3A, Supplementary Fig. 4A**).

**Figure 3.**
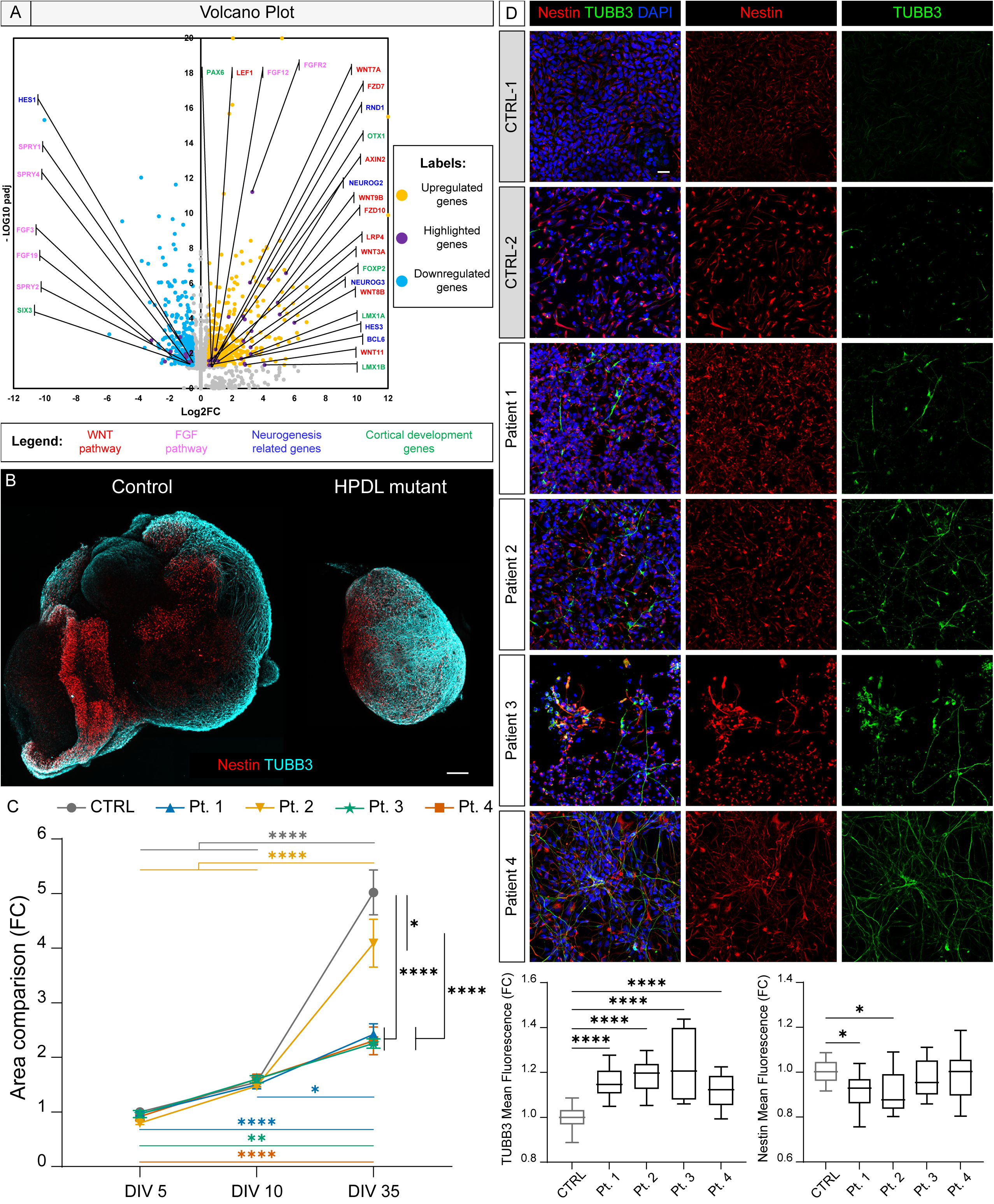
RNA-seq analysis and neurogenesis dysregulation in HPDL neural progenitors. (**A**) Volcano plot showing up- and down-regulated genes in HPDL neural progenitors (DIV16) compared to CTRL counterparts. Highlighted in purple, several DEGs involved in fundamental pathways for cortical development. (**B**) Representative confocal images of Control and HPDL mutant cortical organoids stained for neural progenitor marker Nestin and neuronal specific marker TUBB3. Severe size reduction is striking. Scale bar: 100 μm. (**C**) Organoid growth was followed at different time points and displayed via line plot, showing clear reduction in SPG83-derived cultures (two-way ANOVA, *post-hoc* Tukey’s multiple comparisons test, *n* > 10 organoids from 3 experiments, *P* < 0.0001). All points are represented as mean ± SD. (**D**) Representative confocal images of Nestin and TUBB3 immunostaining in DIV16 CTRL and HPDL cultures, displaying the increase of TUBB3 positive cells in HPDL conditions. Immunofluorescence signals were quantified and represented in box and whisker plots (data are represented as min-to-max). The analysis shows increase of TUBB3 (*n* = 3, *P* < 0.0001) in all patient-derived lines and a decrease in Nestin fluorescence in Patients 1 and 2 compared to CTRLs (*n* = 3, *P* = 0.0212, ordinary one-way ANOVA, *post-hoc* Holm-Šídák’s multiple comparisons test). Scale bar: 25 μm.

To corroborate these data, we differentiated CTRL and SPG83 iPSCs in 3D cortical organoids, following a modified version of an established protocol.^25^ We monitored longitudinally CTRL and HPDL-derived cortical organoid growth at DIV5, 10, and 35. HPDL mutant cortical organoids failed to thrive, showing significant reduced growth compared to controls (**Fig. 3B**). In 3 out of 4 mutant lines, organoids only doubled the size within the analyzed timeframe, whereas CTRL organoids displayed an almost 5-fold increase (**Fig. 3C**). Notably, this phenotype is highly reminiscent of the most severe NEDSWMA-phenotype found in children carrying *HPDL* mutations.^6,7^ Exhaustion of proliferative cortical progenitor pools by means of premature neurogenesis (possibly associated with an increase in apoptosis) constitutes one of the classical pathological mechanisms in models of microcephaly.^32–36^

We also stained control and HPDL mutant adherent (2D) cortical cultures at DIV16 for neural progenitor (Nestin) and neuronal (beta-3 tubulin or TUBB3) specific markers. The majority of the cells differentiated from CTRL lines were Nestin-positive neural progenitors at this stage, with scattered isolated foci of TUBB3-positive differentiated neurons (**Fig. 3D**). Conversely, cortical cultures differentiated from all HPDL lines displayed significant increase of neuronal population, together with reduction of Nestin-positive progenitors in 2 out of 4 cases (**Fig. 3D**), thereby validating occurrence of premature neurogenesis in mutant cells.

Collectively, data coming from different experimental approaches support anticipation of neurogenesis in SPG83 patient-derived cortical cultures.

### HPDL mutant cortical cultures exhibit impairment in deeper layer neurons

Keeping in mind the characteristic “inside-out” pattern typical of neocortical neurogenesis, we performed immunostaining on iPSC-derived cortical neurons (DIV30), expecting enrichment for specific markers of deeper cortical layers (CTIP2 and TBR1, for layers 5 and 6, respectively). Consistent with our previous results, we observed a marked increase in deeper layer neurons in all HPDL mutant lines compared to CTRL (**Fig. 4A**). In particular, increase of CTIP2-positive layer 5 neurons was detected in cortical neurons derived from Patient 2 (16.50 ± 2.04%), Patient 3 (21.94 ± 6.35%), and Patient 4 (56.63 ± 3.10%), compared to the CTRL counterparts (5.69 ± 1.18%; **Fig. 4B**). Interestingly, the increase of layer 5 neurons correlated with a dramatic drop in TBR1-positive layer 6 neurons (0.098 ± 0.029%, 0.21 ± 0.093%, 0.26 ± 0.091% in Patients 2, 3, and 4, respectively, vs 4.14 ± 1.07% in CTRL; **Fig. 4C**). Nevertheless, we observed an increase in the total amount of deeper layer neurons derived from Patients 2, 3, and 4 (**Fig. 4D**). Apparently, the increase in neurogenesis occurred even earlier in cortical cultures from Patient 1, displaying a dramatic increase of layer 6 neurons (9.67 ± 0.67%) and a decrease of layer 5 neurons (1.60 ± 0.43%; **Fig. 4A-C**).

**Figure 4.**
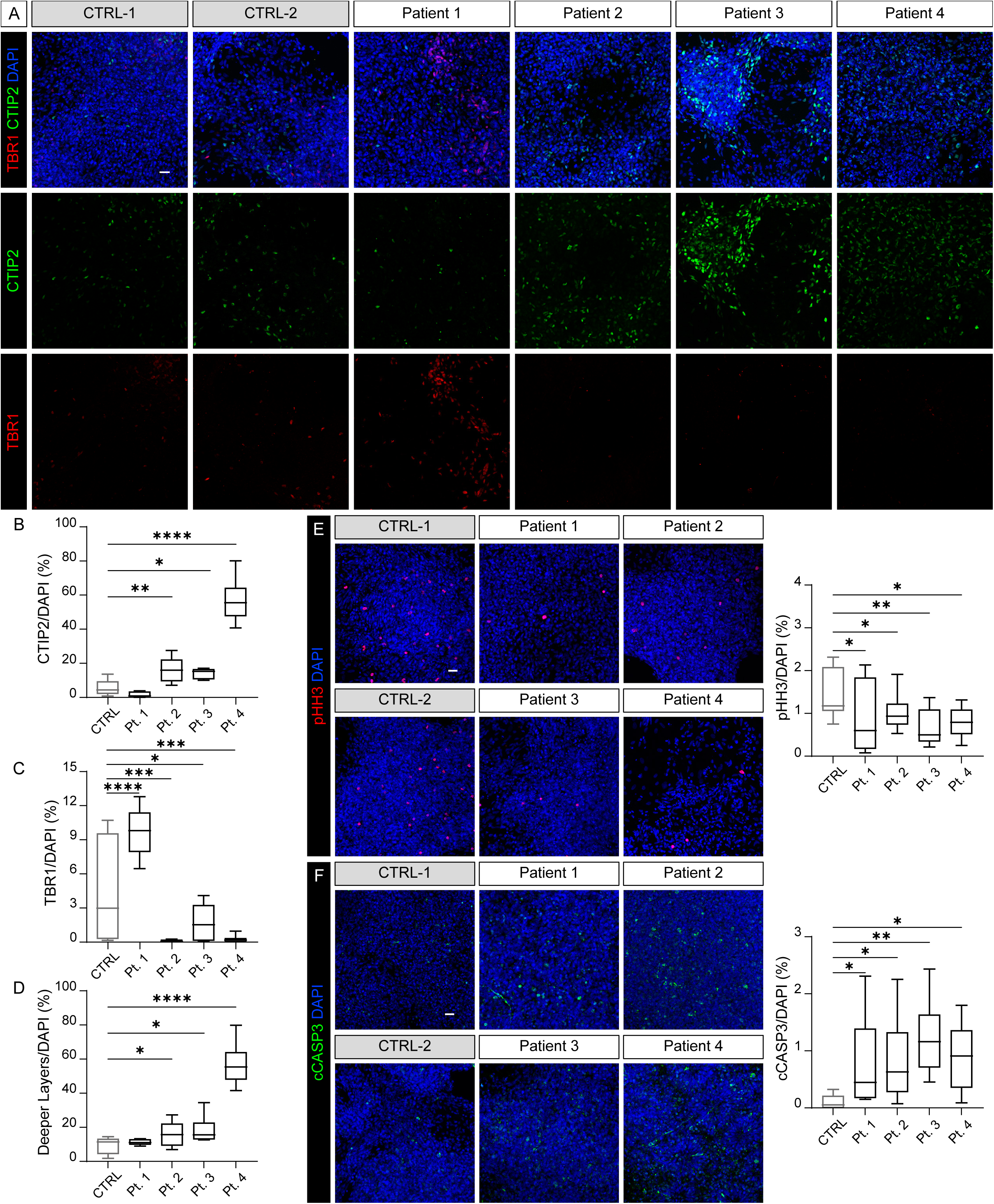
Cortical neurogenesis dysregulation in HPDL neurons. (**A**) Increase in deeper layer neurons in DIV30 SPG83 Patient-derived cortical cultures as shown via immunostaining for layer 5 (CTIP2) and layer 6 (TBR1) cortical markers. Nuclei are stained with DAPI. (**B**) Cell counting on acquired images defined statistically significant increase of CTIP2-positive cells in Patient 2, Patient 3 and Patient 4-derived neurons, while decrease was observed in Patient 1-derived cultures (*n* = 3, *p* < 0.0001). (**C**) TBR1-positive neurons were increased in Patient 1 and decreased in Patients 2, 3, and 4-derived neurons (*n* = 3, *P* < 0.0001). (**D**) Overall quantity of deeper layer (CTIP2+TBR1-positive) neurons results increased all Patients, but not for Patient 1-derived cells (*n* = 3, *P* < 0.0001). (**E**) Reduction of proliferation assessed via positivity for mitotic marker pHH3 in HPDL neurons compared to CTRLs (*n* = 3, *P* = 0.0150). (**F**) Apoptosis is increased in HPDL mutant cultures, as assessed via enhanced positivity for apoptotic marker cCASP3 observed in Patient neurons compared to our two CTRL ones (*n* = 3, *P* = 0.0141). (**A**)-(**F**) All data were represented via box and whisker plots as min-to-max, and all statistic analyses were performed using ordinary one-way ANOVA, *post-hoc* Holm-Šídák’s multiple comparisons test. Scale bars: 20 μm.

Moreover, in line with the observed increase in neurogenesis, HPDL cortical cultures displayed a significant reduction of mitotic cells, positive for phosphorylated Histone H3 (pHH3; **Fig. 4E**). Finally, immunocytochemistry for cleaved Caspase-3 (cCAS3) showed a consistent increase of apoptotic cells at DIV30 in all lines deriving from SPG83 patients (**Fig. 4F**).

These results corroborate the hypothesis that SPG83 patient-derived cortical cultures undergo premature neurogenesis, leading to overall increase of deeper layer neurons combined with a decrease in the number of mitotic cells and an increase of apoptotic cells. Similar findings have been seen in *Hpdl*^-/-^ mouse cerebral cortex.^7^

### Reduction of mitochondrial activity correlates with increase of ROS levels in HPDL mutant cultures

To the end of validating OxPhos defects seen in HPDL KO SH-SY5Y cells, we performed cytochemical staining for COX activity in HPDL DIV30 neurons. Cortical tissues derived from control lines showed an intense and homogeneous signal, coherently with the well-known sustained requirement of oxidative phosphorylation in neuronal cells.^37^ Conversely, SPG83-derived neurons displayed reduced staining (**Fig. 5A**). In keeping with results in SH-SY5Y cells, no functional abnormalities in mitochondrial complex II activity were seen in cultures derived from cell lines analyzed upon SDH cytochemistry (**Supplementary Fig. 4B**).

**Figure 5.**
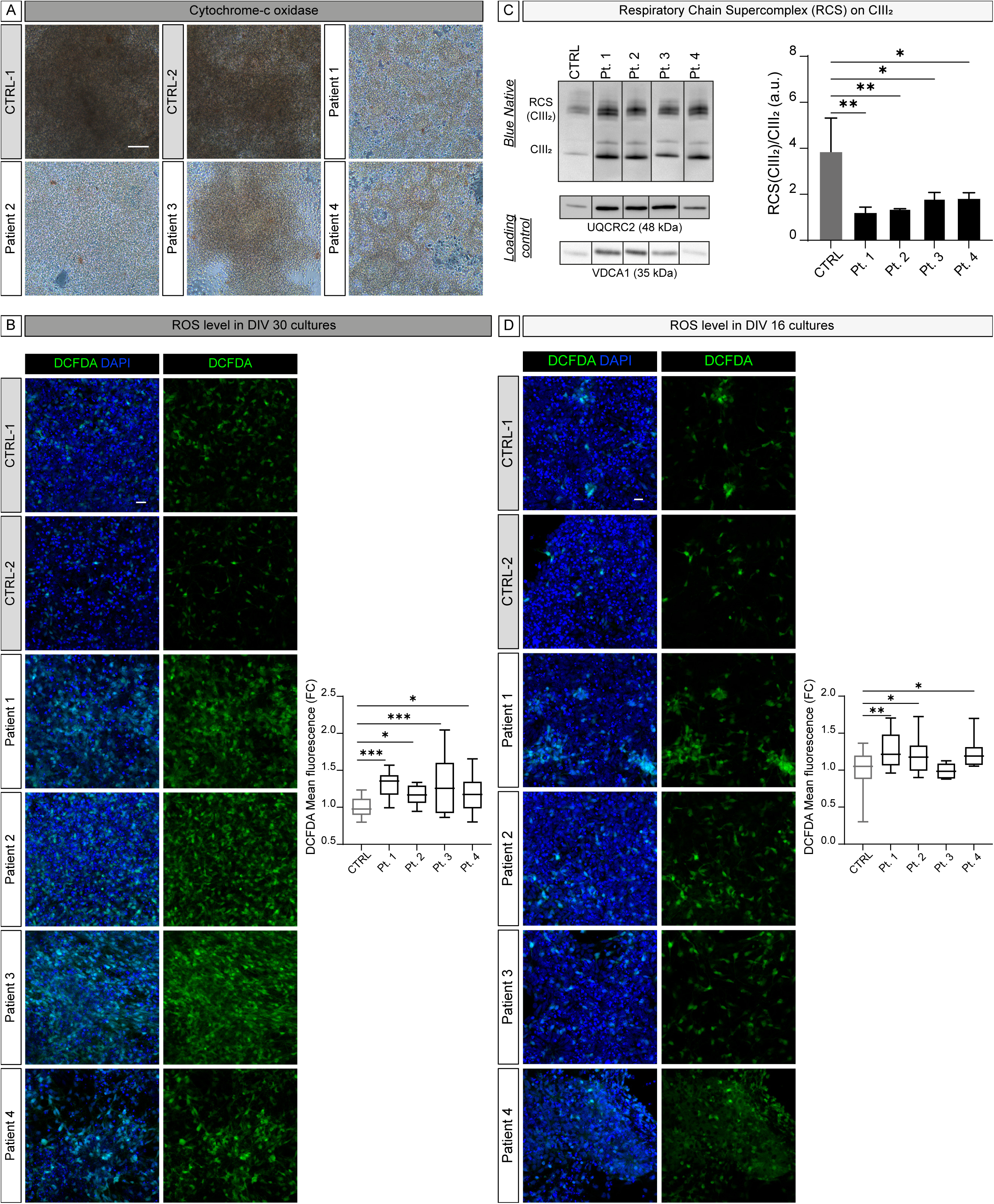
Mitochondrial dysregulation and ROS increase in HPDL cultures. (**A**) SPG83 patient-derived cortical neurons (DIV30) display evident oxidative phosphorylation defects, as shown via Cytochrome-*c* oxidase histochemistry. (**B**) Increase of oxidative stress in DIV30 Patient-derived vs Control neurons, detected via analysis of mean fluorescence of DCFDA probe (in green) and relative quantification depicted in box-and-whisker plot. Enhancement in levels of ROS was significant in all cells carrying HPDL variants (*n* = 3, *P* = 0.0003, ordinary one-way ANOVA, *post-hoc* Holm-Šídák’s multiple comparisons test). (**C**) Representative images of BN-PAGE, loading controls, and corresponding quantifications (in bar plots, represented as mean ± SD) exhibiting the decreased ratio of RCS(CIII_2_) on CIII_2_ in HPDL lines compared to CTRL counterparts (*n* = 3, *P* = 0.0047, ordinary one-way ANOVA, *post-hoc* Holm-Šídák’s multiple comparisons test). (**D**) Representative confocal images of ROS probe DCFDA in DIV16 CTRL and Patient-derived neural progenitors. Quantifications were plotted in box-and-whisker graph (data represented as min-to-max). Statistical analysis proved increase of DCFDA mean fluorescence in cultures derived from Patients 1, 2, and 4 (*n* = 3, *P* = 0.0029, ordinary one-way ANOVA, *post-hoc* Holm-Šídák’s multiple comparisons test). Scale bars are 25 μm in **A** and 20 μm in **B** and **D**.

Since one of the most likely consequences of mitochondrial dysfunction consists in the increase of oxidative stress, we assessed intracellular ROS levels in DIV30 cortical cultures via the cell-permeant dye DCFDA. The experiments showed an actual increase in ROS production in all four mutant lines when compared to control neurons (**Fig. 5B**), confirming high oxidative stress as a putative common hallmark in HPDL disease. Based on these results, we attempted to characterize mitochondrial morpho-functional properties also in cortical progenitors (DIV16), the first stage where we observed a distinctive phenotype in HPDL cortical cultures and investigated the stability of mitochondrial RCSs by BN-PAGE. Similar to KO neuroblastoma cells, DIV16 progenitors derived from SPG83 lines showed both a significantly reduced capacity of complex III to be included in supercomplexes (**Fig. 5C**), and unchanged levels of CoQ_10_, even increased in Patient 1 derived cells (**Supplementary Fig. 4C**).

Finally, mutant DIV16 cells showed increase ROS production (**Fig. 5D**), confirming the link between mitochondrial dysfunction, increase of oxidative stress and regulation of neurogenesis.

### Premature neurogenesis is rescued by antioxidant treatment in HPDL mutant cortical progenitors

Based on previous data, we performed a 1-vs-1 pilot experiment, treating a control (CTRL-1) and a single HPDL mutant line (Patient 2) with two powerful antioxidants, GSH-MEE and MitoTEMPO, a mitochondrial specific ROS scavenger to validate the link between premature neurogenesis in HPDL mutant cortical cultures and a concomitant increase of ROS. In addition, since sustained upregulation of WNT pathway has emerged from RNA-Seq experiments in early mutant cultures, we also tested a known potent WNT inhibitor, XAV939, and assessed the immunostaining signal of TUBB3 to evaluate the levels of neurogenesis (**Fig. 6A**). Whist XAV939 treatment did not modify the phenotype of HPDL lines, a short-term (4 days) administration of each antioxidant compound showed partial rescue of the anticipated neuronal production in HPDL mutant cells, with MitoTEMPO having higher efficiency compared to GSH-MEE (60.85% vs 41.19%; **Fig. 6B**). These results enforce our hypotheses, linking oxidative stress and premature neurogenesis in HPDL neuronal cultures.

**Figure 6.**
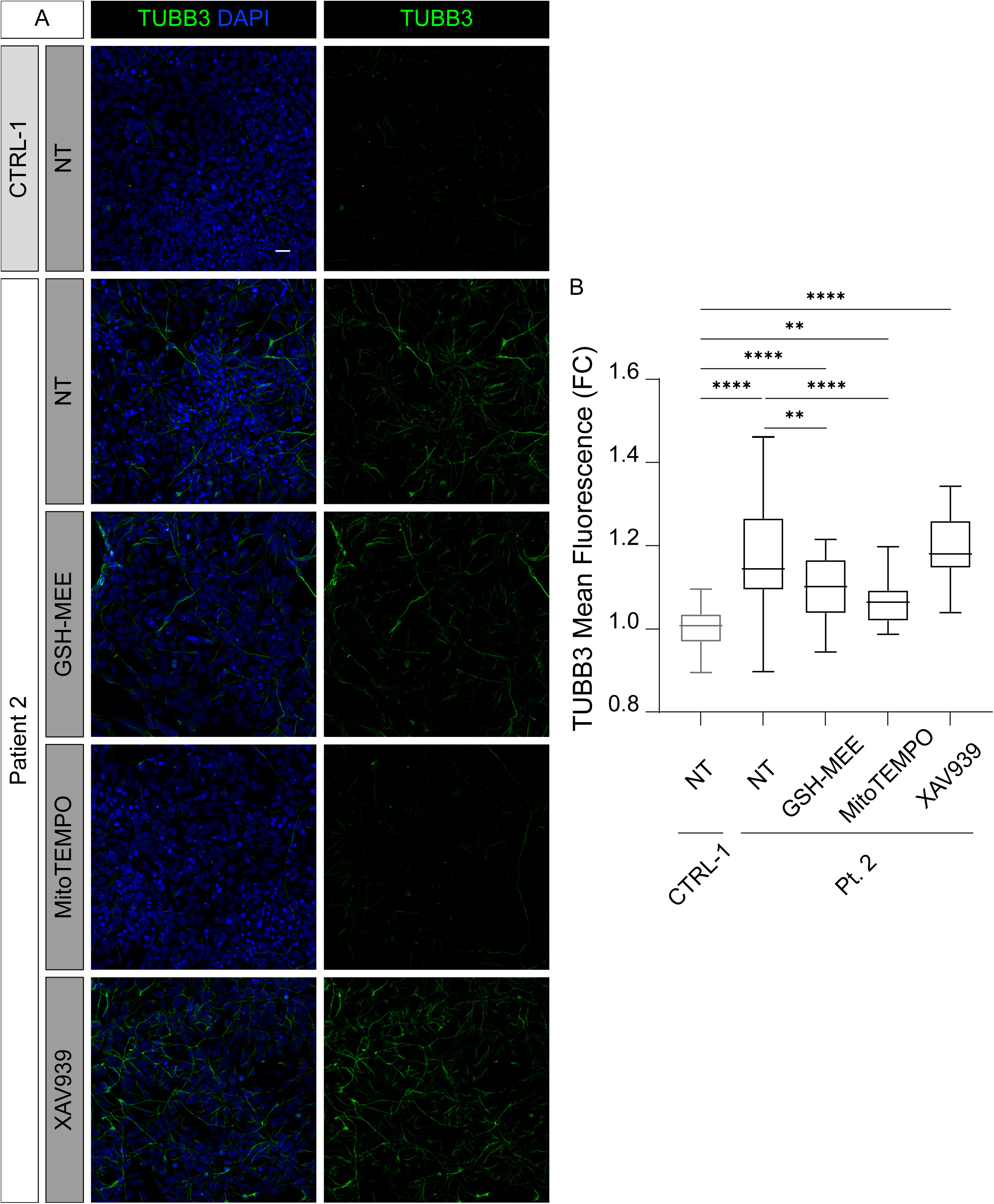
Rescue of increased neurogenesis with antioxidant treatment in SPG83-derived cortical cells. (**A**) Representative confocal images of neuronal marker TUBB3 in CTRL-1 and Patient 2 derived neural progenitors. HPDL cells are not treated (NT) or treated for 4 days with two powerful antioxidants, GSH-MEE or MitoTEMPO, or WNT inhibitor XAV939. Antioxidant administration shows a decrease in TUBB3 positivity compared to NT cells derived from Patient 2. Scale bar: 20 μm. (**B**) TUBB3 mean fluorescence quantification demonstrates a partial rescue after antioxidant treatment, particularly efficient for the case of MitoTEMPO. All data are represented as min-to-max in a box-and-whisker plot; statistical analysis was performed via ordinary one-way ANOVA, *post-hoc* Holm-Šídák’s multiple comparisons test (*n* = 3, *P* < 0.0001)

## Discussion

Human brain development is a finely tuned spatiotemporal-regulated process that can lead to severe neurological disorders if perturbated. In the past few years, the fundamental role of metabolism (and consequently of mitochondrial activity) as neurodevelopmental regulator has increasingly emerged.^38–41^ In this scenario, our findings highlight the role of HPDL in cortical neurogenesis and mitochondrial function. SPG83, a recently identified form of HSP, is caused by mutations in the *HPDL* gene, and characterized by degeneration of corticospinal tract as well as possible presentation of impairments affecting white matter, cerebellar, and cortical structures. *HPDL* encodes a protein of unknown function but deemed to be involved in mitochondrial metabolism and CoQ_10_ biosynthesis. However, its precise role in neural and brain development remains unclear. Our research seeks to address part of this knowledge gap.

We first knocked-out HPDL in neuroblastoma cells and substantiated its role in OxPhos metabolism, observing reduced mitochondrial RC activities and impaired formation of mitochondrial RCS without reduction of CoQ_10_ levels. Importantly, as tumoral cells strongly rely on glycolytic pathways in standard culture conditions, SH-SY5Y cells were kept with limited access to glucose for 48h to enhance OxPhos requirement. As a natural consequence of the mitochondrial dysfunction shown, we observed an increase of ROS levels in conditions of oxidative stress, further corroborating mitochondrial impairment in the absence of HPDL. Transcriptomic analyses showed dysregulated expression of genes involved in axonal development and synapse assembly and organization, strongly suggesting an important role of HPDL in cortical formation and consistent with the impaired neurodevelopment seen in SPG83 children.

To demonstrate the possible impact of HPDL functional ablation in brain development, we extended our investigation to cortical neurons and organoids differentiated from SPG83 patient-derived iPSC lines.^16^ Mutant cell lines successfully achieved cortical specification, with all cultures expressing the telencephalic marker FOXG1 but immunocytochemical characterization of early cultures illustrated premature neurogenesis in SPG83-derived cells already at DIV16. A slight increase in neurogenesis as the one described here could seem almost negligible at a superficial analysis, but the accumulation of such an effect during the highly proliferative phase of cortical development could lead to an increasing depletion of neuroepithelial cells, limiting heavily the expansion of the cortical tissue. The existence of such a scenario is also supported by RNA-seq data in cells at the same early time point of differentiation. Apparently, DIV16 HPDL mutant cells show profound changes in gene expression compared to CTRL counterparts, as revealed by strong upregulation in neurogenesis-related pathways such as NOTCH (with high upregulation of master genes like

NEUROGENIN-2 and -3) and WNT, alongside downregulation of pathways more related to proliferation (as FGF) and increase of genes specific to cortical development. As depletion of proliferative cortical progenitors due to premature neurogenesis represents a well-documented pathological mechanism in microcephaly, we performed *in vitro* 3D-differentiation observing that. SPG83 patient-derived cortical organoids exhibited a reduced growth, reminding the most severe microcephalic cases reported in patients.^6,7^ Despite HPDL mutant iPSCs used in this study derived from donors not presenting severe symptoms, cortical organoid culture represents still an imperfect model, missing many of the non-neural cell populations such as microglia, pericytes, meningeal cells, vascular structures, etc., that actually constitute the final structure of the adult cerebral cortex. The lack of buffering activity provided by these cells, usually exacerbates the phenotypes observed in brain organoids, providing also an advantage for research purpose, as demonstrated by accelerated aging and formation of amyloid and alpha-synuclein aggregates respectively in 3D models of Alzheimer’s^42–46^ and Parkinson’s disease.^47–50^

During neural development, cortical progenitors go through a highly proliferative phase, followed and accompanied by a strictly regulated process of neurogenesis in which newly born neurons populate and form the cortical plate with an “inside-out” organization. Following this model, cortical progenitors change competence of generating different neuronal cell types over time, going from deeper layer (layers V-VI) to upper layer neurons (layer II-III-IV) and finally switching to glia.^35,51^ Consistently with the observed anticipation in neurogenesis, all cell lines deriving from SPG83 patients displayed at a later time point with both an increase of deeper layer neurons and a decrease in proliferating cells suggesting a key role of HPDL to maintain the correct neurodevelopmental timing. Another important modifier in this scenario could be constituted by cell death, physiologically present at basal levels during cortical development and considered to be a distinctive hallmark in numerous models of microcephaly, as it occurs in mice carrying functional ablation in the *Nde1*,^52^ *Lis-1*,^32^ *Mcph1*,^53^ *Aspm*,^54^ and *Diaph3* genes.^33^ Consistent to the phenotype reported in murine *Hpdl* KO cortices,^7^ mutant cell lines displayed raised apoptotic rate in DIV30 cultures. All these data suggest a crucial role for the HPDL protein in the maintenance of cortical progenitor proliferation, regulating the balance between proliferation, neurogenesis, and apoptosis strictly required to build the complexity of the human brain. Notably, lack of Spatacsin, another HSP-related protein, leads to a similar phenotype in cortical organoids cultured from SPG11 patient-derived iPSCs.^55^ Whether a similar role is also played by additional HSP proteins and the detailed mechanisms through which HPDL and Spatacsin regulate neocortex formation remain unclear.

The known mitochondrial localization of HPDL protein, along with the aforementioned results in KO neuroblastoma cells, prompted further investigations in neural progenitors and DIV30 cortical neurons allowing the following considerations.

First, we confirmed HPDL mutant cortical progenitors display a reduction in the assembly of mitochondrial complexes in supercomplexes. As demonstrated in yeast^56^ and zebrafish,^57^ these supramolecular aggregates can boost respiration, increasing maximal respiratory activity and granting sufficient metabolism efficiency (in yeast) to achieve normal body size and fertility (in fish). The functional significance of RCSs is still debated, as mouse models with drastically reduced levels of respirasomes have been described under physiological conditions.^58^ Notably, patients harboring mutations in the complex IV assembly gene *SURF1* present Leigh syndrome^59^ and patient-derived cortical organoids display reduced size and mitochondrial dysfunction reminiscent of what we observe in SPG83.^60^

A second consideration emerging from our study point to increased ROS production as a major player in brain development, as ROS have been described to accumulate in the newt brain and mouse cortical plate from neurogenesis stage onward.^61,62^ DFCDA staining allowed us to deem high oxidative stress as a main hallmark for SPG83 patient-derived cortical cultures, with increase of ROS found even at neuroepithelial stage (DIV16). This set of results suggests the manipulation of ROS as valid therapeutic opportunity not only for SPG83, but also for other genetic forms of HSP, particularly (but not exclusively) those involving mitochondrial proteins, such as SPG7, SPG13, SPG20, SPG31, and SPG74.^1,3^ Compounds with strong antioxidant activity have been used to treat HSP animal models, such as methylene blue and N-acetyl cysteine in SPG4^63^ or the flavonoids naringenin and resveratrol in SPG31.^64,65^ Unfortunately, oral bioavailability and blood brain barrier penetration could constitute a big hurdle for the use in clinical practice. In our case, short time treatment of antioxidants partially rescued the pro-neurogenic effect caused by HPDL ablation. In a translational perspective, the two compounds have a likelihood for attempts *in vivo*. The limitations of this study are multiple including extension to a larger set of patient-derived lines, lack of mechanistic insight on respiratory chain complexes, longer treatments with antioxidants and with different doses, and the use of FDA approved drugs to facilitate translation to clinical practice.

Summarizing, our experiments collectively point to the critical requirement of HPDL protein in neurodevelopment and in the maintenance of mitochondrial morpho-functional stability, supporting cortical progenitor survival and tuning the balance between proliferation and neurogenesis.

The increasingly close bond between neurodevelopment and oxidative metabolism could provide new insights into the pathogenesis of HSPs and other related neurological disorders, potentially offering therapeutic strategies in SPG83 and similar conditions.

## Acknowledgements

We thank Federico Cremisi (Scuola Normale Superiore, Pisa), Paolo Porporato (University of Torino), and all the members of Molecular Medicine Lab for overall theorical and practical scientific support.

## Supplementary material

Supplementary material is available at *Brain* online.

