## Supplementary Materials for "HPDL is critical in human cortical development via regulation of mitochondrial functional properties"

### Supplementary Materials and Methods

#### Generation of CRISPR/Cas9-mediated KO cell lines

The HPDL CRISPR single guide RNA (sgRNA) sequence 5'-CCTCCCGTTCGCGTGCGGGACG-3' was designed in order to efficiently target the *HPDL* gene with a minimal risk of off-target, and the experiment was completed as reported.[1] In particular, SH-SY5Y were transfected with Lipofectamine 2000 (11668019, Thermo Fisher Scientific) and selected with puromycin. Genotype of resistant clones was performed by Sanger sequencing. Tracking of Indels by Decomposition (SYNTHEGO) web tool (https://ice.synthego.com/#/), was used to accurately characterize and quantify the induced genome editing events. One SH-SY5Y HPDL stable KO clone was selected for downstream analyses.

#### Western Blot

For Western blot (WB) analysis, samples were homogenized in M-PER™ mammalian protein extraction reagent (78501, Thermo Fisher Scientific) containing inhibitors of proteases (78429, Thermo Fisher Scientific); immunoassays were performed following described procedures.[2] Antibodies used were anti-HPDL (1:1000, HPA031997, Sigma-Aldrich; specificity verified in Sun et al.[3]), anti-β-tubulin (1:1000, 2146, Cell Signaling Technology). The secondary antibodies used were peroxidase-conjugated anti-rabbit (1:5000, 111-035-045, Jackson Labs).

#### Determination of reactive oxygen species in neuroblastoma cells

Intracellular reactive oxygen species (ROS) production was assessed using a cellular detection kit (ab113851, Abcam). Cells were incubated with 25 μM 2′,7′-dichloro-dihydrofluorescein diacetate ($H_2$DCFDA) for 45 minutes at 37°C. Afterwards, cells were cultured for an additional hour in the presence/absence of 150 μM tert-butyl hydrogen peroxide (TBHP), a ROS-inducing compound. ROS levels were then measured using a SpectraMax® ID3 plate reader (Molecular Devices) at excitation and emission wavelengths of 485 nm and 535 nm, respectively. The fluorescence data, expressed as relative fluorescent units (RFU), were adjusted for background signal and normalized to the Hoechst 33342 intensity (H3570, Thermo Fisher Scientific) for each well.

#### Immunostaining, cell counting, and mean fluorescence analysis

Different cell types were treated in the same way, fixing with 4% formaldehyde (157-4-100, Electron Microscopy Science) for 12 min at RT and permeabilizing in PBS with 0.5% Triton X-100 (X100, Sigma) for 10 min at RT. PBS with 5% Normal Goat Serum (NGS; S-1000, Vector Laboratories) and

0.3% Triton X-100 was used as blocking solution for 1 h at RT and incubated with primary antibodies (FOXG1, 1:500, ab18259, Abcam; TUBB3, 1:200, ab41489, Abcam; CTIP2, 1:500, ab18465, Abcam; TBR1, 1:500, ab31940, Abcam; pHH3 1:500, 9706, Cell Signaling; cCASP3, 1:500, 9661, Cell Signaling; OCT4, 1:1000, ab19857, Abcam; NANOG 1:200, 4903, Cell Signaling; SSEA4 1:400, 4755, Cell Signaling; TRA1-60, 1:1000, 4746, Cell Signaling; Brachyury, 1:500, 81694S, Cell Signaling; Nestin, 1:500, ab18102, Abcam; SOX17, 1:500, 81778s, Cell Signaling) in antibody solution (PBS with 3% NGS and 0.2% Triton X-100) at 4 °C overnight. Cells were then washed 3 times with PBS and incubated in the same antibody solution with secondary antibodies (Goat anti-rabbit IgG (H+L) 555, 1:500, a21429, Invitrogen; Goat anti-mouse IgG (H+L) 647, 1:500, a21236, Invitrogen; Goat anti-chicken IgG (H+L) 647, 1:500, a32933, Invitrogen; Goat anti-mouse IgG (H+L) 488, 1:500, a11029, Invitrogen; Goat anti-rat IgG (H+L) 488, 1:500, a11006, Invitrogen) for 1 h at RT. All nuclei were counterstained with DAPI (1 μg/ml; 10236276001, Merck). All images were acquired with a Zeiss LSM 900 confocal microscope (Zeiss) and processed with Fiji software (ImageJ 1.54f). Cell counts were executed by blinded operators via Cell Counter plugin (Fiji) and normalized on total number of nuclei.

#### Lipid Extraction and HPLC Analysis

Cells were cultured for 48 h with OxPhos medium, harvested cells and resuspended in 0.2 mL of ice-cold PBS. To each sample, 4 μl of $CoQ_4$ μM were added to a final concentration of 2 μM, as an internal standard to control the efficiency of extraction. Then, 600 μl of ice-cold extraction solution (ethanol:hexane, 1:2) were added. The tubes were vortexed for 10 min and then centrifuged at 13˙200 g for 5 min at 4°C. The upper phase was collected and put into an 1 ml amber glass mass spectrometry vial. The samples were dried under nitrogen flow at 30°C and then were resuspended in 100 μl of mobile phase (0.1 M lithium perchlorate, 80% acetonitrile, 20% methanol). Lipid extracts were subjected to HPLC analysis for measurement of $CoQ_{10}$ levels. About 10 μl of sample was injected and separated with a Waters Alliance e2695 separation module under a flow rate of 0.25 ml/min at 45°C. The column used for the analysis was XBridge C18 3.5 μm 1.0x100 mm (186003127, Waters). The CoQ signal was registered with the Waters 2489 UV/Visible Detector at 275 nm. The chromatogram of each sample was analyzed evaluating the area under the peak of $CoQ_4$ (internal standard) and $CoQ_{10}$ (endogenous CoQ species). A standard curve for $CoQ_{10}$ in the range 0.1 to 4 μM with $CoQ_4$ at 4μM was constructed for absolute quantification of $CoQ_{10}$ normalized to protein content.

#### Mitochondrial purification and Blue Native-PAGE

Prior to the measurements, cells were cultured for 48 h with OxPhos medium. Blue Native-PAGE (BN-PAGE) followed by immunoblotting was performed to identify respiratory chain complexes bands. Mitochondrial enriched fractions were obtained following the protocol outline in Frezza et al.[4] About 35 μg of mitochondrial proteins (4 g/g digitonin/protein ratio) were loaded in BN-PAGE gels and run first for 30 minutes at 150 V with Cathode Buffer Dark Blue, and then for 120 minutes at 150 V with Cathode Buffer light Blue. Mitochondrial complexes were transferred to a PVDF membrane. WB was performed following standard procedures and using the following antibodies: anti-NDUFB8 (1:1000, 4592210, Invitrogen), anti-SDHA (70KDa Fp subunit) (1:1000, 459200, Invitrogen), anti-Cox-II (MTCO2 Monoclonal Antibody 12C4F12) (1:1000, A6404, Invitrogen), anti-UQCRC2 (1:1000, ab14745, Abcam), and anti-ATP5A (1:1000, ab14748, Abcam). The secondary antibody used was peroxidase-conjugated goat anti-mouse (1:2000, 32430, Thermo Scientific). Images were acquired after the chemiluminescence reaction (LiteAblot Turbo, Euroclone) by the iBright FL1500 (Agilent) instrument. All original gels are shown in Supplementary Fig. 7. Quantifications of single bands were performed via GelAnalyzer 19.1 software.

#### Measurement of Oxygen Consumption by the Seahorse XFe96 Extracellular Flux Analyzer

The oxygen consumption rate (OCR) was determined using a Seahorse XFe96 Extracellular Flux Analyzer (Agilent) following the manufacturer's instructions. XF96 V3 PS cell culture microplates (103792-100, Agilent Technologies) were pre-coated with 0.1 mg/ml poly-D-lysine and then 2.5 x $10^5$ cells for each well were seeded. Cells were seeded in OxPhos medium and incubated for 24 hrs. Prior to the measurements, the medium was replaced with Seahorse XF base medium supplemented with 2 mM glucose, 2 mM glutamine, and 1 mM sodium pyruvate and incubated for 1 h at 37°C without $CO_2$. XF cell mito stress test protocol was run according to manufacturer’s instructions. OCR was measured under basal conditions and after sequential addition of oligomycin 2 μM, FCCP 1.5 μM (C2920, Merck), rotenone 0.5 μM (R8875, Merck), and antimycin A 0.5 μM (A8674, Merck). Respiration rates were normalized to μg/ml of protein.

#### mtDNA quantification

Mitochondrial DNA (mtDNA) quantity was determined by Real time PCR from 0.4 to 10 ng of total DNA extracted from SH-SY5Y cells by the kit Puregene Cell and tissue kit (158043, Qiagen). The analysis was performed using the Rotor-Gene 6000 Corbett Research instrument, the SensiFAST™ SYBR® No-ROX kit (BIO-98005, Meridian), and primers for the mitochondrial gene *MT-ND1* (F:

CCCTAAAACCCGCCACATCT, R: GAGCGATGGTGAGAGCTAAGGT) and the nuclear gene *FAS* (F: GGCTCTGTGAGGGATATAAAGACA, R: CAAACCACCCGAGCAACTAATCT). The relative quantification of mtDNA *vs* nuclear DNA was performed by the software of the instrument following the method of $^{\Delta\Delta}$Ct.

#### Transcriptomics analyses

RNA quality control was performed on 48 RNA samples. RNA integrity was assessed using the RNA 6000 Nano Kit on a Bioanalyzer (Agilent Technologies). RNA samples were quantified using the Qubit RNA BR Assay Kit on a Qubit fluorometer (Thermo Fisher Scientific). All samples met the requirements for RNA sequencing. Illumina RNA-seq libraries were generated using the TruSeq stranded mRNA ligation kit (Illumina) from 400 ng of RNA samples, after poly(A) capture and according to manufacturer's instructions. Quality and size of RNAseq libraries were assessed by capillary electrophoretic analysis with an Agilent 4150 Tape station (Agilent). Libraries were quantified by real-time PCR against a standard curve with the KAPA Library Quantification Kit (KapaBiosystems, Wilmington, MA, USA). Illumina Sequencing - Libraries were pooled at equimolar concentration and sequenced in 150PE on a NovaSeq6000 (Illumina) generating on average 29.8 million fragments per sample.

Quality of reads was assessed using FastQC software (http://www.bioinformatics.babraham.ac.uk/projects/fastqc/). Raw reads were trimmed with fastp (v.0.21.0)[5] with --trim_poly_x parameter, to remove adapters and low-quality bases with default parameters. Filtered reads were aligned to the Homo sapiens reference genome (Ensemble version 109) using STAR aligner (v2.7.9a) with parameter --peOverlapNbasesMin 5. Reads distribution on CDS, intronic and intergenic was computed using reads_distribution.py script from RSeQC suite (v4.0.0).[6] Gene expression quantification has been performed using RSEM and the Homo sapiens Ensembl v.111 annotation on the two different time-points and over-time. Genes-level abundance estimated counts and gene length obtained with RSEM (v1.3.1; https://github.com/deweylab/RSEM) were summarized into a matrix using the R package tximport (within DESeq2 package) and subsequently the differential expression analysis was performed with DESeq2 v.1.38.3.[7] To generate more accurate log2 fold change estimates for low expressed genes, the shrinkage of the Log2 FoldChange was performed applying the apeglm method.[8] For transcriptomics of neuroblastoma cells, Gene Ontology (GO) enrichment analysis of DEGs with padj< 0.05 and LFC <|1| have been performed using ClusterProfiler (v3.18.1) (Guangchuang Yu, et al., OMICS 2012). Subsequently, the top 30 enriched Biological Processes GO categories have been plotted. For transcriptomic analysis of patient-derived cells at DIV16, GO enrichment and gene clustering for up- and downregulated categories was performed via STRING,[9] keeping for both padj< 0.05 and LFC <|0.5|. Chord and bar plots were generated using SRplot.[10]

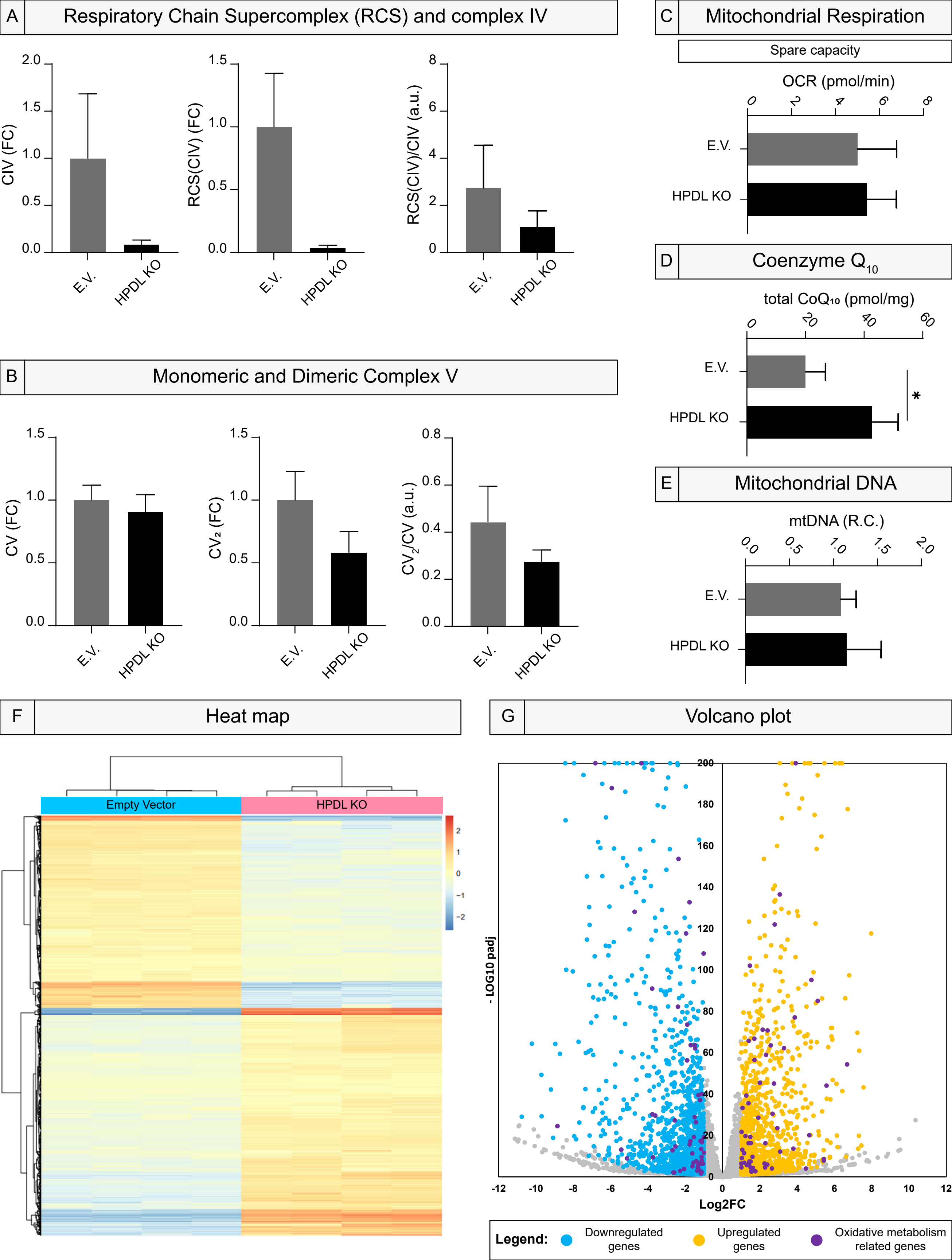
A Respiratory Chain Supercomplex (RCS) and complex IV
CIV (FC)
RCS(CIV) (FC)
RCS(CIV)/CIV (a.u.)
E.V.
HPDL KO
B Monomeric and Dimeric Complex V
CV (FC)
CV2 (FC)
CV2/CV (a.u.)
C Mitochondrial Respiration
Spare capacity
OCR (pmol/min)
D Coenzyme Q10
total CoQ10 (pmol/mg)
*
E Mitochondrial DNA
mtDNA (R.C.)
F Heat map
Empty Vector
HPDL KO
G Volcano plot
- LOG10 padj
Log2FC
Legend:
Downregulated genes
Upregulated genes
Oxidative metabolism related genes

**Supplementary Figure 1. RCS and complex IV, dimeric complex V, mitochondrial analysis, total Coenzyme $Q_{10}$ quantification, and RNA-Seq in HPDL KO SH-SY5Y cell line.** (**A**) Bar plots show a tendency to decrease of CIV, RCS(CIV), and RCS(CIV)/CIV in HPDL KO line compared to E.V. one (unpaired Student's t-test with Welch's correction, $n$ = 3, $P$ = 0.1461; unpaired Student's t-test with Welch's correction, $n$ = 3, $P$ = 0.0596; unpaired Student's t-test, $n$ = 3, $P$ = 0.2135, respectively). All data are represented as mean ± SD. (**B**) Graphs exhibit the relative quantification of monomeric and dimeric complex V (CV and $CV_2$, respectively) and the ration between them ($n$ = 3, $P$ = 0.4271; $n$ = 3, $P$ = 0.0635; $n$ = 3, $P$ = 0.1413). All data are represented as mean ± SD. All statistical analyses were performed via unpaired Student's t-test. (**C**) Seahorse XF analysis demonstrates no differences in spare capacity for HPDL KO compared to E.V. SH-SY5Y cell lines. All data in bar plots are represented as mean ± SD, unpaired Student's t-test ($n$ = 3, $P$ = 0.7529). (**D**) Bar plot shows the increased amount of total $CoQ_{10}$ levels in HPDL KO compared to E.V. cells. All data are represented as mean ± SD, unpaired Student's t-test ($n$ = 3, $P$ = 0.0254). (**E**) Bar plot indicating no difference between mtDNA abundance between HPDL KO and E.V. cell lines. Data are represented as mean ± SD($n$ = 3, $P$ = 0.7691, unpaired Student's t-test). (**F**) Heat map derived from RNA-seq on E.V. and HPDL KO cells, clearly represents a binary expression of genes. (**G**) Volcano plot showing dysregulated expression of genes in HPDL KO compared to E.V. cells. In evidence (purple), genes involved in oxidative metabolism.

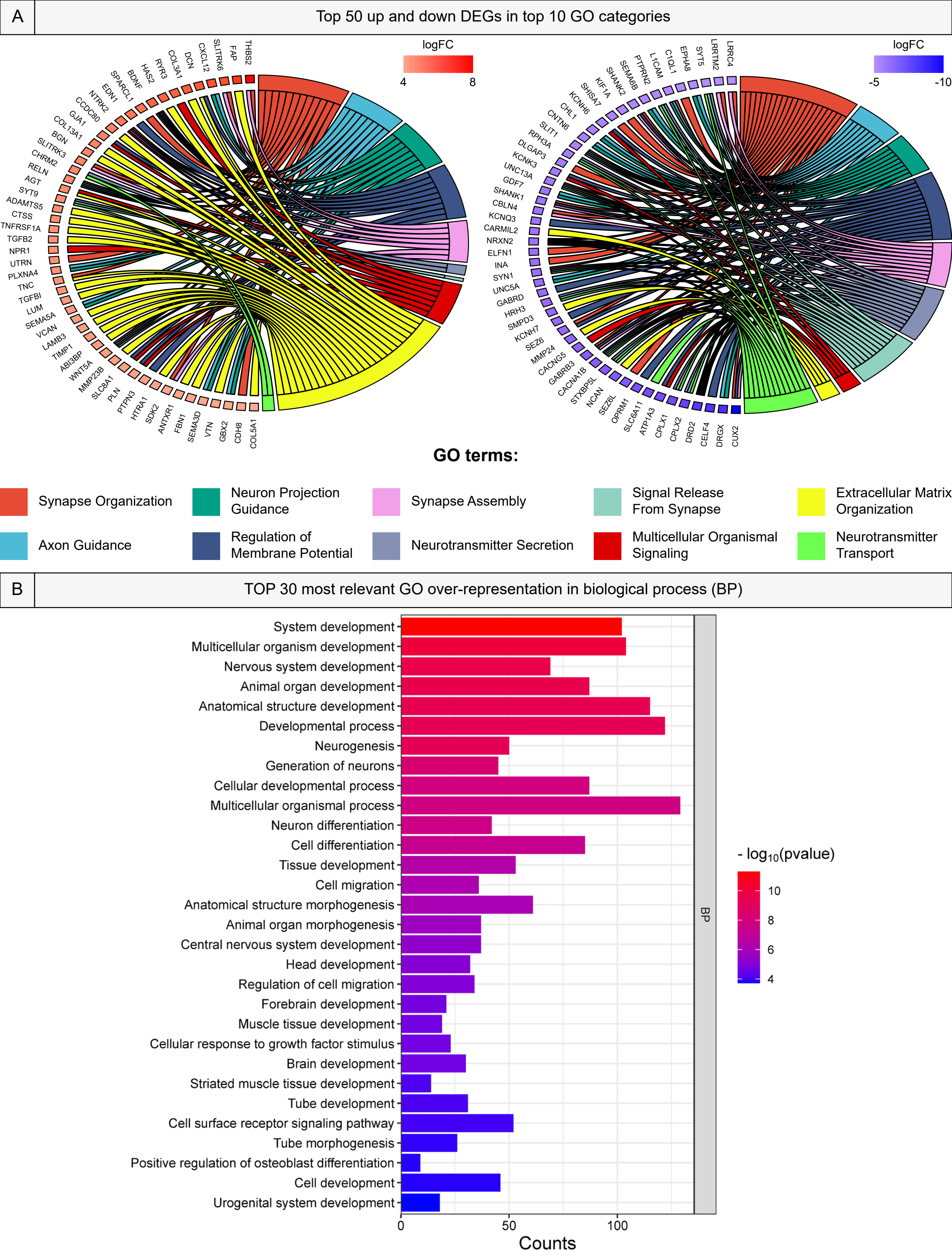
A
Top 50 up and down DEGs in top 10 GO categories
logFC
4
8
THBS2
FAP
SLITRK6
CXCL12
DCN
COL3A1
RYR3
HAS2
BDNF
SPARCL1
EDN1
NTRK2
CCDC80
GJA1
COL13A1
BGN
SLITRK3
CHRM2
RELN
AGT
SYT9
ADAMTS5
CTSS
TNFRSF1A
TGFB2
NPR1
UTRN
PLXNA4
TNC
TGFBI
LUM
SEMA5A
VCAN
LAMB3
TIMP1
ABI3BP
WNT5A
MMP23B
SLC8A1
PLN
PTPN3
HTRA1
SDK2
ANTXR1
FBN1
SEMA3D
VTN
GBX2
CDH8
COL5A1
logFC
-5
-10
LRRC4
LRRTM2
SYT5
EPHA8
C1QL1
L1CAM
PTPRN2
SEMA6B
SHANK2
KIF1A
SHISA7
KCNH6
CHL1
CNTN6
SLIT1
RPH3A
DLGAP3
KCNK3
UNC13A
GDF7
SHANK1
CBLN4
KCNQ3
CARMIL2
NRXN2
ELFN1
INA
SYN1
UNC5A
GABRD
HRH3
SMPD3
KCNH7
SEZ6
MMP24
CACNG5
GABRB3
CACNA1B
STXBP5L
NCAN
SEZ6L
OPRM1
SLC6A11
ATP1A3
CPLX1
CPLX2
DRD2
CELF4
DRGX
CUX2
GO terms:
Synapse Organization
Neuron Projection Guidance
Synapse Assembly
Signal Release From Synapse
Extracellular Matrix Organization
Axon Guidance
Regulation of Membrane Potential
Neurotransmitter Secretion
Multicellular Organismal Signaling
Neurotransmitter Transport
B
TOP 30 most relevant GO over-representation in biological process (BP)
System development
Multicellular organism development
Nervous system development
Animal organ development
Anatomical structure development
Developmental process
Neurogenesis
Generation of neurons
Cellular developmental process
Multicellular organismal process
Neuron differentiation
Cell differentiation
Tissue development
Cell migration
Anatomical structure morphogenesis
Animal organ morphogenesis
Central nervous system development
Head development
Regulation of cell migration
Forebrain development
Muscle tissue development
Cellular response to growth factor stimulus
Brain development
Striated muscle tissue development
Tube development
Cell surface receptor signaling pathway
Tube morphogenesis
Positive regulation of osteoblast differentiation
Cell development
Urogenital system development
BP
0
50
100
Counts
- log10(pvalue)
10
8
6
4

**Supplementary Figure 2. Chord plots and GO categories in HPDL KO SH-SY5Y cells.** (**A**) Chord plots show TOP 50 up- and downregulated DEGs in TOP 10 GO categories. (**B**) GO analysis exhibits the most relevant significantly different BP categories in HPDL KO vs E.V. cells. Bar plot depicts level of significance (*P*-value, expressed as $-\log_{10}$) and the number of gene counts.

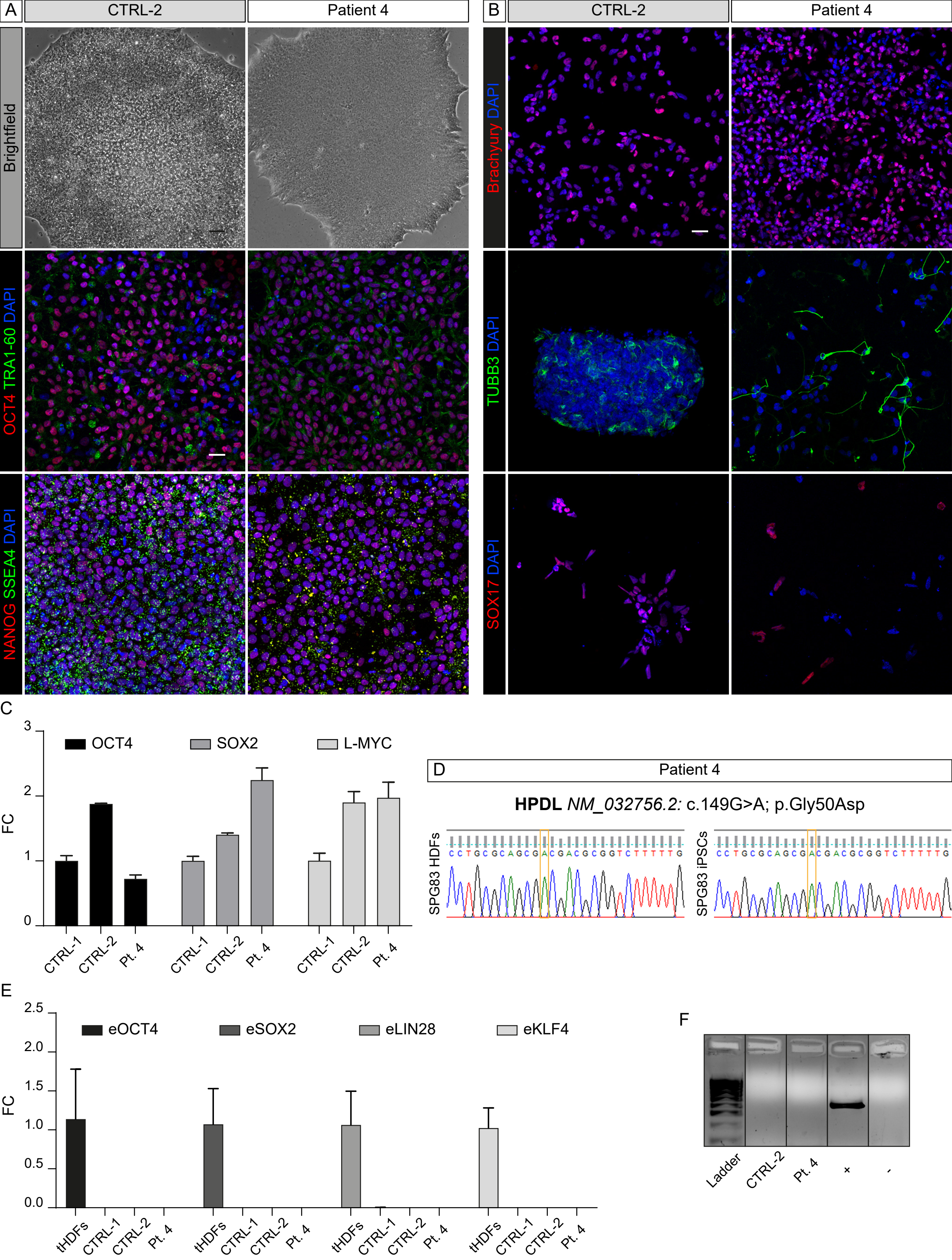

A
CTRL-2
Patient 4
Brightfield
OCT4 TRA1-60 DAPI
NANOG SSEA4 DAPI
B
CTRL-2
Patient 4
Brachyury DAPI
TUBB3 DAPI
SOX17 DAPI
C
OCT4
SOX2
L-MYC
FC
0
1
2
3
CTRL-1
CTRL-2
Pt. 4
D
Patient 4
HPDL NM_032756.2: c.149G>A; p.Gly50Asp
SPG83 HDFs
SPG83 iPSCs
CCTGCGCAGCGACGACGCGGTCTTTTTG
CCTGCGCAGCGACGACGCGGTCTTTTTG
E
eOCT4
eSOX2
eLIN28
eKLF4
FC
0.0
0.5
1.0
1.5
2.0
2.5
tHDFs
CTRL-1
CTRL-2
Pt. 4
F
Ladder
CTRL-2
Pt. 4
+
-

**Supplementary Figure 3. Generation and characterization of hiPSC lines induced from CTRL-2 and Patient 4 fibroblasts.** (**A**) Representative brightfield images showing clone morphology of CTRL-2 and Patient 4 iPSCs (scale bar: 50 µm), and representative confocal images of the same cells stained for stemness markers OCT4, TRA-1-60, NANOG, and SSEA4 (scale bar: 20 µm). (**B**) Representative confocal images of mesodermal (Brachyury), ectodermal (TUBB3), and endodermal (SOX17) markers in differentiated cells derived from all iPSC lines. Scale bar: 20 µm. (**C**) Bar plots show the same or higher expression of OCT4, SOX2, and L-MYC genes in our generated iPSCs compared with gold-standard CTRL-1. (**D**) Electropherograms indicate that the c.149G>A mutation is present in both Patient 4 parental HDFs and iPSCs. (**E**) Bar plots showing the total absence of eOCT4, eSOX2, eLIN28, and eKLF4 episomal gene expression in CTRL-1 (taken as standard) and our iPSCs compared with tHDFs. (**F**) Gel image showing lack of mycoplasma contamination in CTRL-2 and Patient 4-derived iPSCs.

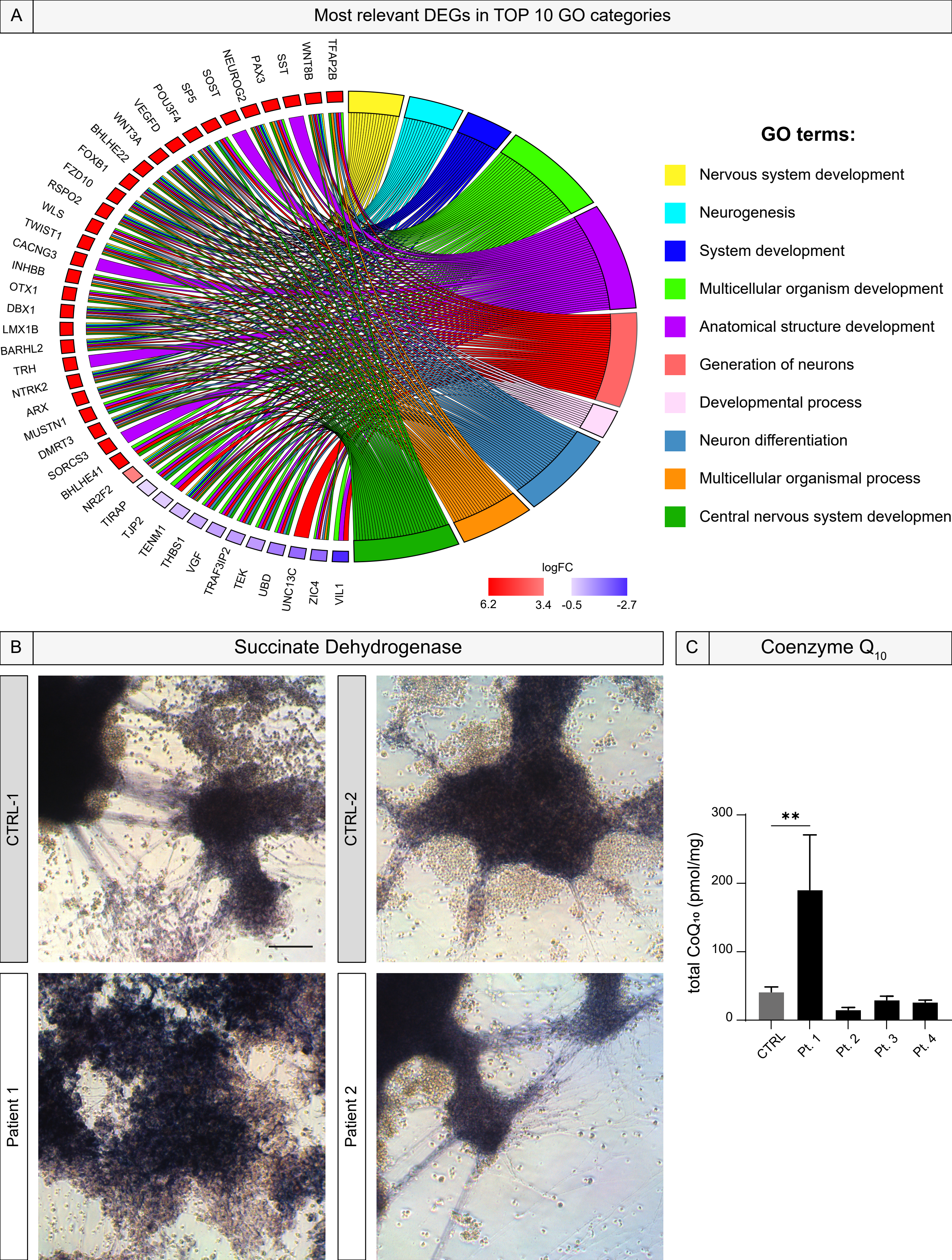
A
Most relevant DEGs in TOP 10 GO categories
TFAP2B
WNT8B
SST
PAX3
NEUROG2
SOST
SP5
POU3F4
VEGFD
WNT3A
BHLHE22
FOXB1
FZD10
RSPO2
WLS
TWIST1
CACNG3
INHBB
OTX1
DBX1
LMX1B
BARHL2
TRH
NTRK2
ARX
MUSTN1
DMRT3
SORCS3
BHLHE41
NR2F2
TIRAP
TJP2
TENM1
THBS1
VGF
TRAF3IP2
TEK
UBD
UNC13C
ZIC4
VIL1
GO terms:
Nervous system development
Neurogenesis
System development
Multicellular organism development
Anatomical structure development
Generation of neurons
Developmental process
Neuron differentiation
Multicellular organismal process
Central nervous system developmen
logFC
6.2
3.4
-0.5
-2.7
B
Succinate Dehydrogenase
CTRL-1
CTRL-2
Patient 1
Patient 2
C
Coenzyme Q10
total CoQ10 (pmol/mg)
300
200
100
0
**
CTRL
Pt. 1
Pt. 2
Pt. 3
Pt. 4

**Supplementary Figure 4. Chord plot from DIV16 RNA-seq, SDH cytochemistry in HPDL cortical neurons, and total Coenzyme $Q_{10}$ quantification in neural progenitors.** (**A**) Chord plot representing up- (TOP 30) and down-regulated DEGs in top 10 GO categories in SPG83 patient derived cortical progenitor cells (DIV16). To be noticed, most categories are related to neurogenesis, and most genes are upregulated. (**B**) SDH cytochemistry indicates that Succinate Dehydrogenase or complex II normally works in HDPL lines, as observed in HPDL KO SH-SY5Y cells. (**C**) Bar plot shows the only increased amount of $CoQ_{10}$ oxidated protein in Patient 1 neural progenitors compared to CTRLs. All data are represented as mean ± SD, ordinary one-way ANOVA, *post-hoc* Holm-Šídák's multiple comparisons test ($n = 3$, $P = 0.0092$).

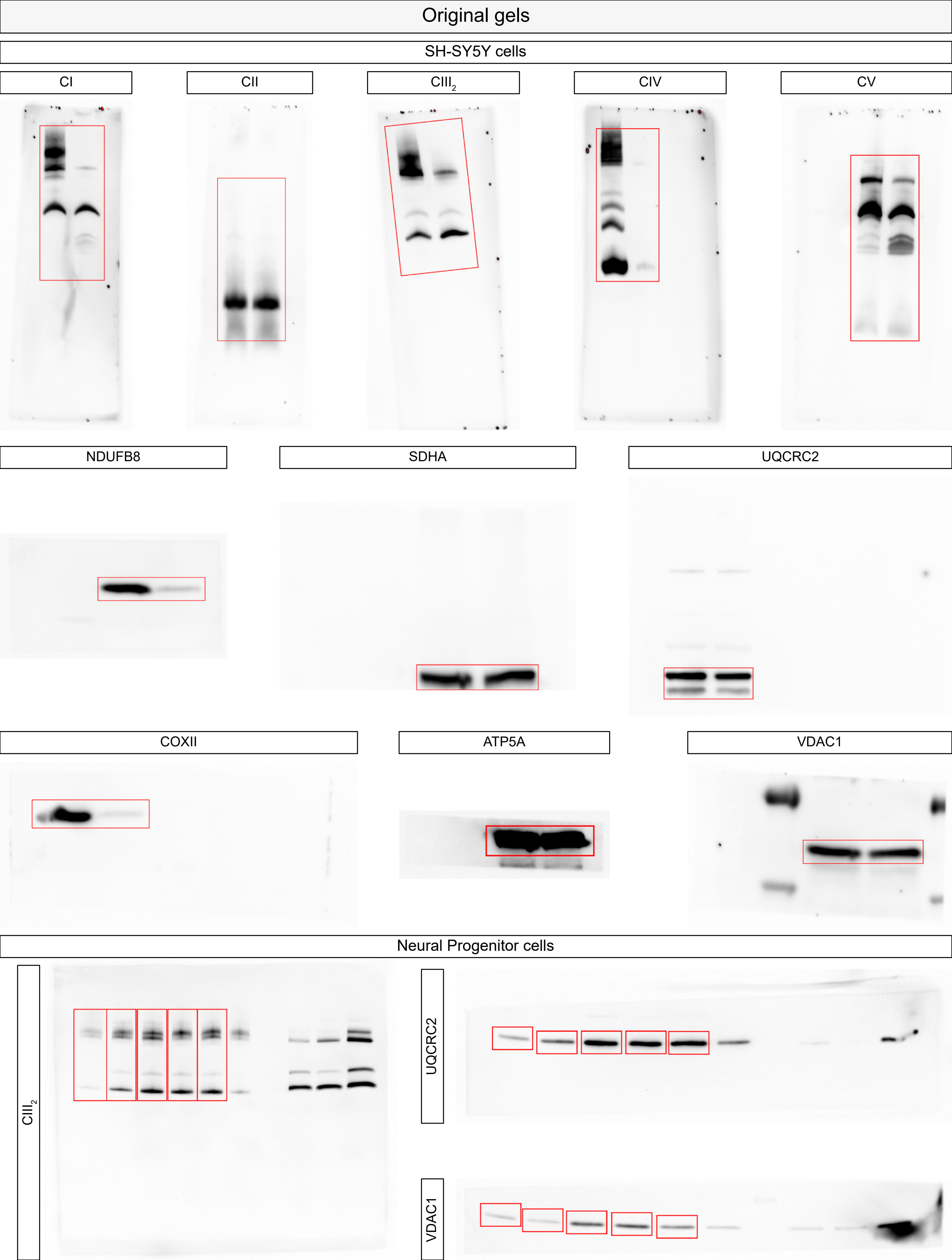
Original gels
SH-SY5Y cells
CI
CII
CIII2
CIV
CV
NDUFB8
SDHA
UQCRC2
COXII
ATP5A
VDAC1
Neural Progenitor cells
CIII2
UQCRC2
VDAC1

**Supplementary Figure 5. Original blots**
